# MLKL deficiency protects against low-grade, sterile inflammation in aged mice

**DOI:** 10.1101/2022.10.17.512454

**Authors:** Emma C. Tovey Crutchfield, Sarah E. Garnish, Jessica Day, Holly Anderton, Shene Chiou, Anne Hempel, Cathrine Hall, Komal M. Patel, Pradnya Gangatirkar, Connie S.N. Li Wai Suen, Alexandra L. Garnham, Andrew J. Kueh, Ueli Nachbur, Andre L. Samson, James M. Murphy, Joanne M. Hildebrand

## Abstract

MLKL and RIPK3 are the core signaling proteins of the inflammatory cell death pathway, necroptosis, which is a known mediator and modifier of human disease. Necroptosis has been implicated in the progression of disease in almost every physiological system and recent reports suggest a role for necroptosis in aging. Here we present the first comprehensive analysis of age-related histopathological and immunological phenotypes in a cohort of *Mlkl*^*-/-*^ and *Ripk3*^*-/-*^ mice on a congenic C57BL/6J genetic background. We show that genetic deletion of *Mlkl*, but not *Ripk3*, in female mice interrupts immune system aging, specifically delaying the age-related reduction of circulating lymphocytes. *Mlkl*^*-/-*^ female mice were also protected against age-related, low-grade chronic sterile inflammation, with a reduced number of inflammatory infiltrates present in the connective and muscle tissue at 17 months relative to wild-type littermates. These observations implicate MLKL in age-related sterile inflammation, suggesting a possible application for long-term anti-necroptotic therapy in humans.

## INTRODUCTION

Necroptosis is a programmed form of lytic cell death that culminates in the rupture of biological membranes and subsequent release of inflammatory, intracellular constituents termed DAMPs (damage-associated molecular patterns)^1^. Nearby sensors of the innate immune system detect DAMP release into the surrounding extracellular space and an inflammatory response ensues^2^. Necroptosis is triggered by a range of both host- and pathogen-derived stimuli, culminating in the phosphorylation of mixed lineage kinase domain-like (MLKL) by the receptor-interacting serine/threonine protein kinase 3 (RIPK3)^3, 4, 5^. Phosphorylated MLKL oligomerises and dissociates from RIPK3, before translocating to biological membranes where it accumulates into higher-order complexes that disrupt plasma membrane integrity^6, 7, 8, 9, 10, 11^.

In mice, the genetic deletion of *Mlkl* or *Ripk3* does not impose overt developmental or early-life homeostatic defects on mice^5, 12, 13, 14, 15^, except for some asymptomatic renovascular^16^, neurovascular^17^ and trabecular^18^ differences reported in *Ripk3*-deficient mice generated on a C57BL/6N background. At 3 months of age, *Mlkl*^*–/–*^ and *Ripk3*^*–/–*^ mice are macroscopically indistinguishable and share similar peripheral blood cell and platelet populations to their wild-type littermates^5, 13, 14, 15, 19^. Yet in response to challenge, *Mlkl*^*–/–*^ and *Ripk3*^*–/–*^ mice often respond differently to their wild-type counterparts, and each other. This has been demonstrated in models of *Staphylococcus aureus* infection^20, 21^, renal ischemia-reperfusion injury^17, 22^, dermatitis^23, 24^ and many others (recently reviewed in ref.^25^). The differences between *Mlkl*^*-/-*^ and *Ripk3*^*-/-*^ mice have been attributed to the necroptosis-independent roles of MLKL and RIPK3, for example in NF-κB activation, inflammasome activation, apoptosis signaling, and endosome/exosome formation^26, 27, 28, 29, 30, 31^. It is important to note, however, that comparisons between *Mlkl*^*–/–*^ and *Ripk3*^*-/-*^ mice are not always performed in parallel and therefore interfacility variability in housing conditions and ambient microbiomes are likely to contribute to divergent phenotypes between studies. Additionally, the most commonly used *Mlkl*^*-/-*^ mouse lines were generated and are maintained on a C57BL/6J background^5, 13^, while the most used *Ripk3*^*-/-*^ mouse line is C57BL/6NJ^15^. Despite being derived from the same original mouse strain, 6J and 6NJ are separated by over 60 years of independent breeding and often demonstrate profound differences in morbidity and mortality following inflammatory challenges, for example, in LPS- or TNFα-induced sepsis^32^ and Influenza A virus infection^33^.

The absence of *Mlkl* and *Ripk3* has been variably reported to influence aging of the male reproductive system in mice. Male *Mlkl*^*-/-*^ and *Ripk3*^*-/-*^ mice were shown to exhibit youthful seminiferous tubules and extended reproductive longevity relative to wild-type males of the same age in an initial study^12^. However, these findings were challenged in a subsequent study that reported no evidence for genotype-dependent testicular aging^34^. Modeling the impact of *Mlkl* and *Ripk3* deficiency in aged laboratory mice will not only inform our understanding of human male reproductive aging, but also offers insights into the effects of long-term use of RIPK3 or MLKL-blocking therapies on aging across all human physiological systems.

As a result of the high prevalence of dysregulated necroptosis in a vast range of human disease^22, 35, 36, 37, 38, 39, 40, 41, 42^, the therapeutic targeting of the necroptotic pathway is being actively pursued on several fronts. While upstream serine/threonine kinase, RIPK1, has been the target of many clinical trials for autoimmune conditions (e.g., rheumatoid arthritis^43^), neurodegeneration (e.g., amyotrophic lateral sclerosis^44^) and cancer (e.g., pancreatic ductal adenocarcinoma^45^), no MLKL- or RIPK3-binding compounds have been included in human trials to date. In light of three recent clinical reports linking *MLKL* loss-of-function mutations to neurodegeneration^46, 47^ and diabetes^48^, modeling the impact of prolonged MLKL loss in mice is integral to understand if therapeutic inhibition of MLKL could contribute to the development or progression of human disease.

Accordingly, here we present a comprehensive characterization of aged *Mlkl*^*-/-*^ and *Ripk3*^*-/-*^ mice relative to wild-type littermate controls. Both strains have been bred, housed, and examined in the same facility under identical conditions, and the *Ripk3*^*-/-*^ knockout strain, published for the first time here, was generated on the C57BL/6J background. These studies have identified a novel role for MLKL in aging, revealing a delayed age-related reduction in circulating lymphocytes and a chronic, low-grade sterile inflammation in female *Mlkl*^*-/-*^, but not *Ripk3*^*-/-*^, aged mice.

## RESULTS

### Generation of *Ripk3*^*-/-*^ mice on a C57BL/6J background

To permit direct comparison with the previously generated *Mlkl*^*-/-*^ C57BL/6J strain^5^, we developed a novel *Ripk3*^*-/-*^ strain on a C57BL/6J genetic background via CRISPR-Cas9 mediated disruption of the mouse *Ripk3* locus **(Supplementary Figure 1A)**. Mice were born according to the Mendelian ratio and showed no overt signs of disease, consistent with the previously described *Ripk3*^*-/-*^ strain that was generated using homologous recombination on a C57BL/6NJ background^15^ **(Supplementary Figure 1B)**. Immunoblots of protein extracts from a range of organs demonstrated that RIPK3 was readily detectable in all examined organs in wild-type mice – heart, colon, ileum, spleen, testes, and liver – but was absent in tissues of *Ripk3*^*-/-*^ mice, verifying deletion of the *Ripk3* protein product **(Supplementary Figure 1C)**. Immortalised mouse dermal fibroblasts (MDFs) isolated from wild-type and *Ripk3*^*-/-*^ mice were examined for their sensitivity to death stimuli. As expected, *Ripk3*^*-/-*^ MDFs did not die in response to TSI-induced (TNF; Smac-mimetic, Compound A; pan-caspase inhibitor, IDN-6556; termed TSI) necroptosis, with the hallmark of the necroptosis pathway activation – the phosphorylation at Serine 345 within mouse MLKL^5^ – not observed **(Supplementary Figure 1D, E)**. In response to TNF-induced apoptosis (TNF and Smac-mimetic Compound A; termed TS), *Ripk3*^*-/-*^ MDFs exhibited reduced levels and kinetics of death in comparison to wild-type MDFs **(Supplementary Figure 1E)**. These data validate successful RIPK3 deletion in a mouse strain of C57BL/6J background, which can be used as a tool to compare phenotypic differences between otherwise congenic *Mlkl*^*-/-*^ and *Ripk3*^*-/-*^ mouse strains.

### *Mlkl*^*-/-*^ and *Ripk3*^*-/-*^ mice show no overt evidence of age-related reproductive system degeneration

To determine the long-term consequences resulting from necroptotic signaling deficiency, a cohort of wild-type littermate-controlled *Mlkl*^*-/-*^ and *Ripk3*^*-/-*^ mice were aged to 12 months and extensively phenotyped. As observed previously, 12-month-old *Mlkl*^*-/-*^ and *Ripk3*^*-/-*^ mice displayed no overt disease or change in rates of mortality^12, 14, 34^. In contrast to previous reports, body weights of male *Mlkl*^*-/-*^ mice, and male and female *Ripk3*^*-/-*^ mice were, on average, equivalent to wild-type littermate controls^12, 16^ **(Figure 1A)**. Female *Mlkl*^*-/-*^ mice were, however, on average, 12% heavier than wild-type littermate controls **(Figure 1A)**. This increase in body weight was not driven by differences in the weight of any one specific organ we measured **(Figure 1B – D, Supplementary Figure 1F – H)**. The weights of all organs were comparable between *Mlkl*^*-/-*^ or *Ripk3*^*-/-*^ and their respective wild-type littermate controls **(Figure 1B – D, Supplementary Figure 1F – H**) with the exception of the spleen in *Ripk3*^*-/-*^ males. When presented as a percentage of body weight, *Ripk3*^*-/-*^ males had a 12% decrease in relative spleen weight compared to wild-type littermate controls, however the splenic architecture remained intact and no macroscopic pathologies were observed **(Figure 1B, E**). In support of recent observations^34^, 12-month-old *Mlkl*^*-/-*^ and *Ripk3*^*-/-*^ males had on average, equivalent seminal vesicle and testes weights compared to their respective wild-type littermate controls **(Figure 1C, D**). Overall, the absence of *Mlkl* and *Ripk3* at 12 months of age did not result in any overt pathogenic phenotypes in the major reproductive and non-reproductive organs.

**Figure 1.**
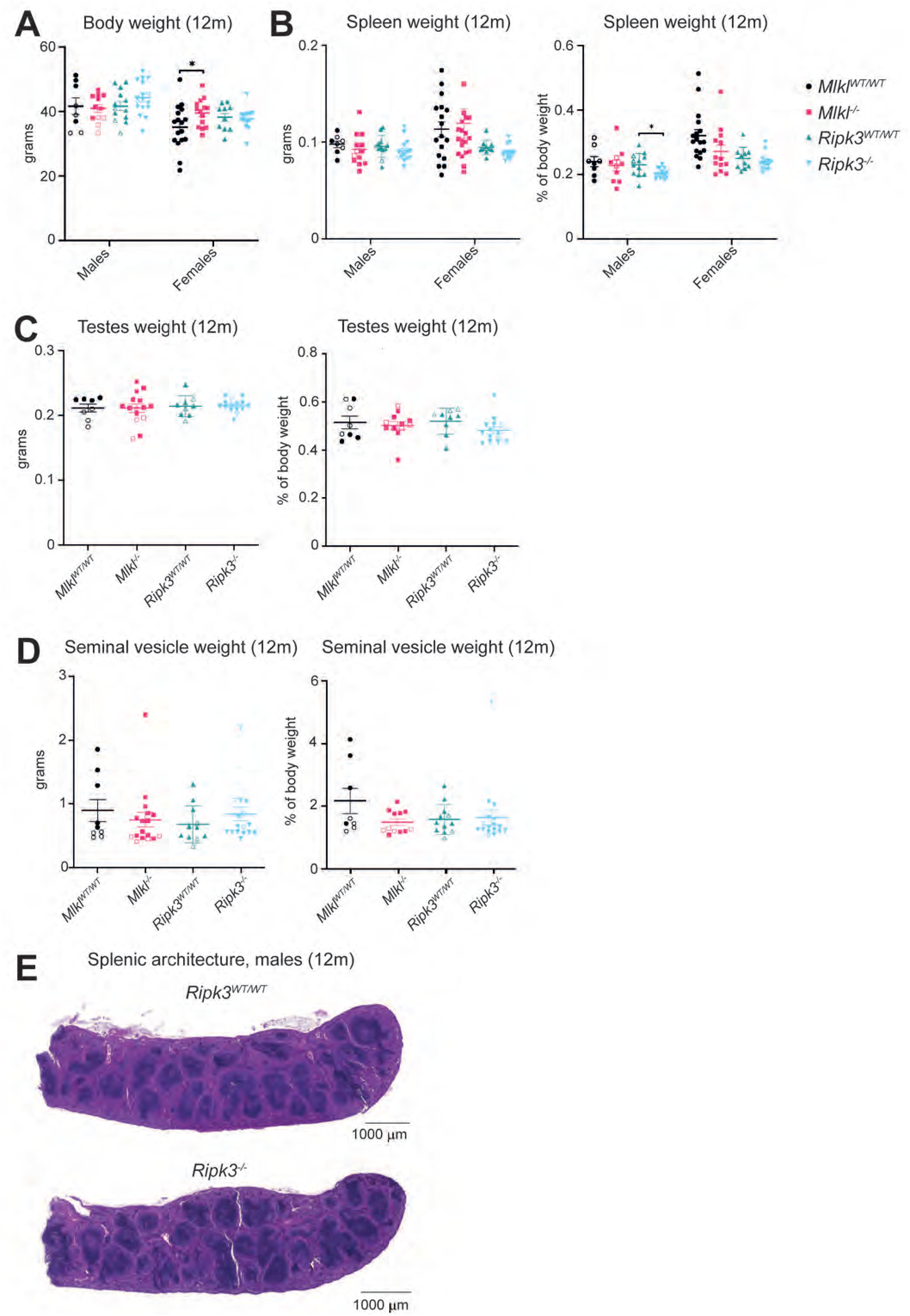
*Ripk3*^*-/-*^ male mice exhibit reduced spleen weight in comparison to wild-type littermate controls. **(A)** Body weights of *Mlkl*^*-/-*^, *Ripk3*^*-/-*^ and wild-type littermate control mice at 12 months of age. Weight (absolute and relative) of 12-month-old *Mlkl*^*-/-*^, *Ripk3*^*-/-*^ and wild-type littermate control spleen **(B**), testes **(C**) and seminal vesicles **(D). (E)** H&E-stained section of male *Ripk3*^*-/-*^ mice and wild-type littermate control at 12 months of age. Each symbol represents a datum from one individual mouse and error bars represent mean ± SEM for *n* = 8 – 20. Hollowed-out symbols represent data from fighting male mice. H&E-stained sections of the spleen **(E)** are representative of *n* = 3 mice per genotype. \**p* < 0.05 was calculated using an unpaired, two-tailed Student’s t-test.

### *Mlkl*^*-/-*^ mice do not develop overt neurodegenerative or diabetic disease at 9 months of age

In response to three recent clinical reports of human *MLKL* loss-of-function mutations associated with severe neurological or diabetic disease^46, 47, 48^, aged mice deficient in *Mlkl* or *Ripk3* were assessed for signs of these conditions. The patients identified in these reports presented with disease onset as late as 30 years of age therefore, mice were aged to 9 months which roughly equates to 35 human years^49^. These mice were examined by professional histopathologists, blind to genotype. All mice, irrespective of genotype, were classified as ostensibly normal and displayed intact gross motor and neurological function as measured by gait and the tail suspension test^50^ **(Supplementary Figure 2A, Supplementary File 1**). None of the characterised neurodegenerative lesions present in male human carriers of homozygous *MLKL* loss-of-function mutation, *p*.*Asp369GlufsTer22* (rs56189347) were observed in brain sections of knockout and wild-type littermate control mice^47^ (**Figure 2A, Supplementary Figure 2B**). For example, there was no evidence of global atrophy or periventricular white matter lesions observed in H&E-stained brain sections, and full body X-rays showed no evidence of osteoporosis, such as bone thinning or fractures, which are common comorbidities associated with neurological disorders^51^ (**Figure 2A, B, Supplementary Figure 2B, C, Supplementary File 1**). Furthermore, all brains were symmetrical and similar in size, measured by length, width, and height (**Supplementary Figure 2D**). A rare missense loss-of-function mutation in *MLKL* was also found to exclusively segregate with the affected male and female siblings suffering from maturity-onset diabetes of the young (MODY)^48^. Random blood glucose measurements taken from *Mlkl*^*-/-*^ and *Ripk3*^*-/-*^ mice at 6 and 12 months revealed no pre-diabetic or diabetic phenotypes. At both 6 and 12 months of age, *Mlkl*^*-/-*^ and *Ripk3*^*-/-*^ mice had equivalent random blood glucose measurements relative to littermate controls **(Figure 2C, D)**. Further, H&E-stained pancreas sections of 9-month-old mice revealed no lesions of significance, with typical exocrine and endocrine tissue architecture **(Figure 2E, F)**.

**Figure 2.**
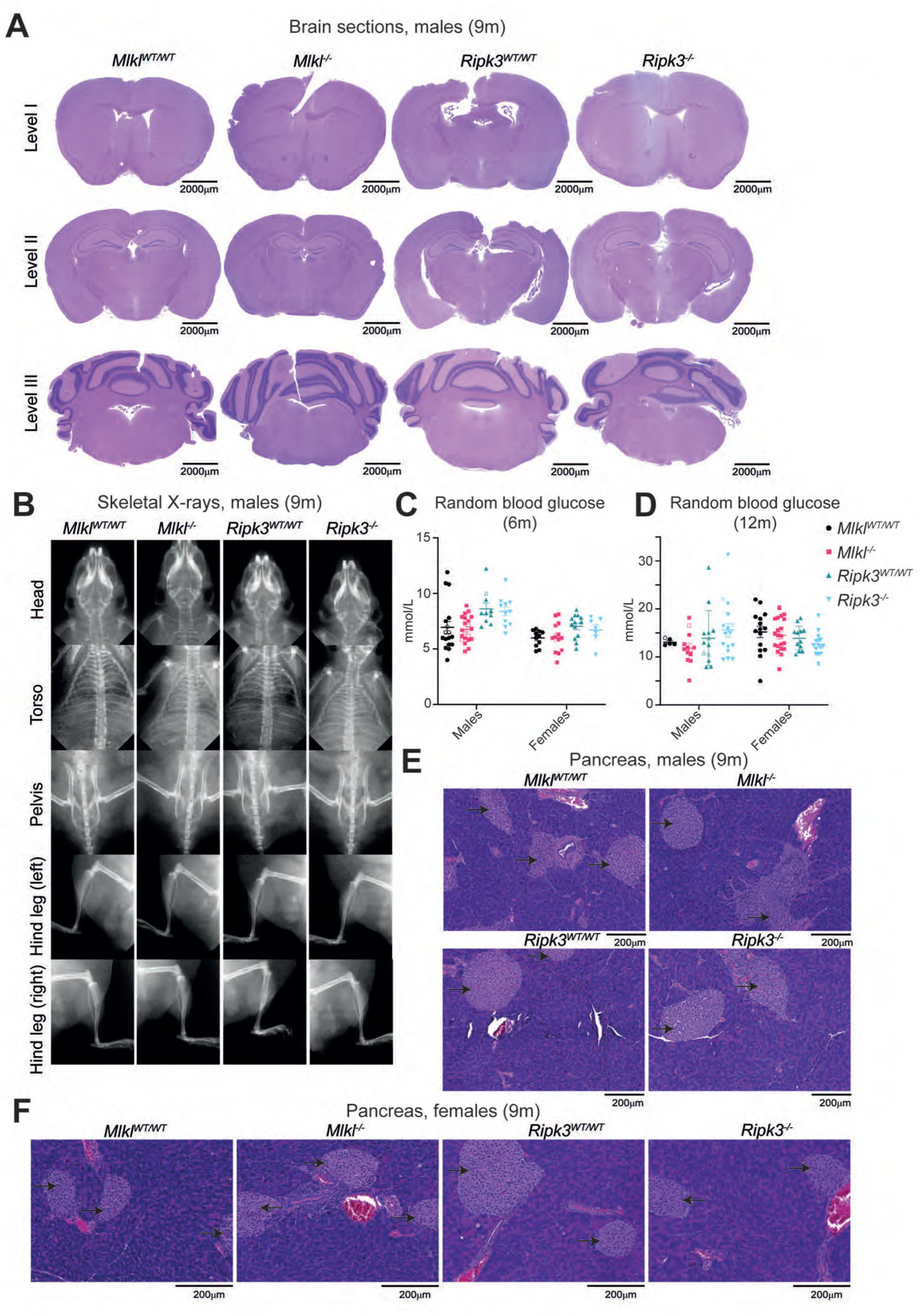
No overt macroscopic signs of neurodegenerative or diabetic disease were observed in 9-month-old *Mlkl*^*-/-*^ and *Ripk3*^*-/-*^ mice. **(A)** H&E-stained sections of the levels I, II and III of the brain and **(B)** X-ray images of the head, torso, pelvis, and hind leg from 9-month-old male *Mlkl*^*-/-*^, *Ripk3*^*-/-*^ and wild-type littermate control mice. Level I includes the cortex, corpus callosum, caudate putamen and lateral ventricles; level II includes the hippocampus, thalamus, hypothalamus, and lateral and third ventricles, and level III, the cerebellum, pons and fourth ventricle. Random blood glucose measurements for 6-**(C)** and 12-month-old **(D)** *Mlkl*^*-/-*^, *Ripk3*^*-/-*^ and wild-type littermate control mice. H&E-stained sections from the pancreas of 9-month-old male **(E)** and female **(F)** *Mlkl*^*-/-*^, *Ripk3*^*-/-*^ and wild-type littermate control mice. Islets of Langerhans are denoted by black arrows. H&E-stained sections **(A, E, F)** and X-ray images **(B)** are representative of *n* = 2 – 5 mice per genotype. Each symbol represents a datum from one individual mouse and error bars represent mean ± SEM for *n* = 5 – 21. Hollowed-out symbols represent data from fighting male mice. \**p* < 0.05 and \*\**p* < 0.01 were calculated using an unpaired, two-tailed Student’s t-test.

### *Mlkl*^*-/-*^ mice exhibit age-dependent increases in peripheral lymphocyte numbers

Extending the phenotyping of our aged *Mlkl* and *Ripk3* deficient cohort, peripheral blood cell numbers were quantified in *Mlkl*^*-/-*^ and *Ripk3*^*-/-*^ mice from 3 – 14 months. Fourteen-month-old male and 12-month-old female *Mlkl*^*-/-*^ mice displayed, respectively, a 57% and 44% increase in the average number of peripheral white blood cells (WBC) compared to their wild-type littermate controls **(Figure 3A)**. This change in WBC number was driven by increases in the peripheral lymphocyte population. In comparison to wild-type littermate controls, peripheral lymphocytes were elevated by 61% in 14-month-old male *Mlkl*^*-/-*^ mice and by 51% in 12-month-old female *Mlkl*^*-/-*^ mice **(Figure 3B)**. Three-month-old female *Mlkl*^*-/-*^ mice also displayed a 26% increase in the average number of peripheral WBC in comparison to wild-type littermate controls **(Figure 3A)**. This change, however, was reflected in a 53% increase in the average number of peripheral neutrophils in comparison to wild-type littermate controls **(Figure 3C)**. Similarly, *Mlkl*^*-/-*^ male mice demonstrated an 81% increase in the average number of peripheral neutrophils and yet a 58% decrease in the average number of peripheral monocytes at 3 months compared to wild-type littermate controls **(Figure 3C, D)**. By 9 months of age, male *Mlkl*^*-/-*^ mice exhibited a contrasting 60% reduction in their average neutrophil numbers compared with wild-type littermate controls **(Figure 3C)**. Despite these significant variations in white blood cell counts, bone marrow and peripheral blood smears from 9- and 17-month-old *Mlkl*^*-/-*^ mice demonstrate morphologically indistinguishable lymphoid and myeloid cells to wild-type littermate controls **(Supplementary File 1)**.

**Figure 3.**
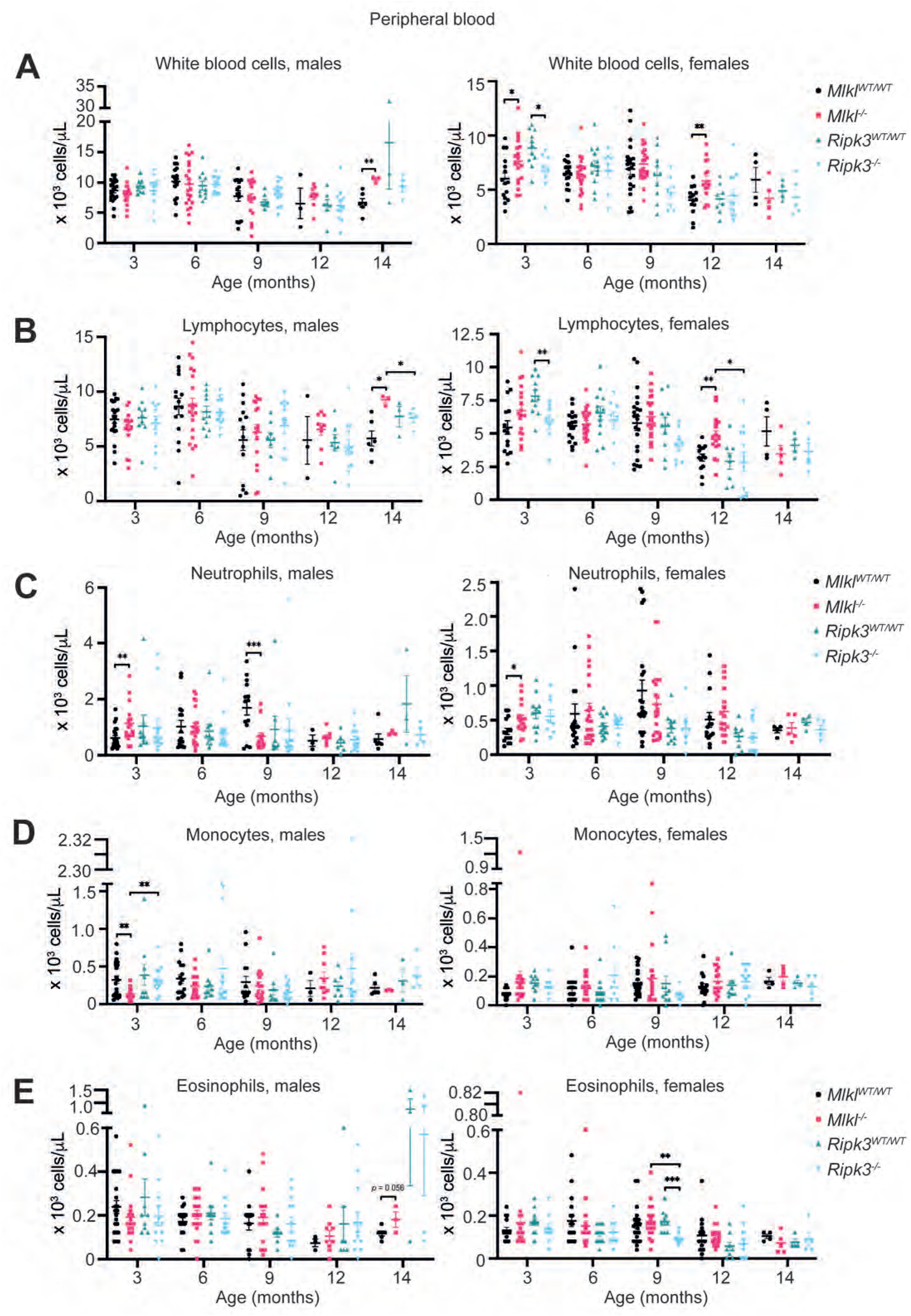
Male and female *Mlkl*^*-/-*^ mice exhibit increased circulating lymphocytes with age. ADVIA hematology quantification of circulating white blood cells **(A)**, lymphocytes **(B)**, neutrophils **(C)**, monocytes **(D)** and eosinophils **(E)** in sex separated *Mlkl*^*-/-*^, *Ripk3*^*-/-*^ and wild-type littermate control mice from 3 – 14 months of age. Each symbol represents a datum from one individual mouse and error bars represent mean ± SEM for *n* = 3 – 22. \**p* < 0.05, \*\**p* < 0.01, \*\*\**p*< 0.001 calculated using an unpaired, two-tailed Student’s t-test.

Female, but not male, *Ripk3*^*-/-*^ mice also exhibited significant differences in their peripheral WBC populations across age **(Figure 3A – E)**. A 24% reduction in the mean circulating WBC population was observed in female *Ripk3*^*-/-*^ mice compared to wild-type controls at 3 months **(Figure 3A)**. Again, this difference was reflected in a 25% decrease in the average number of peripheral lymphocytes compared to wild-type littermate controls **(Figure 3B)**. Female *Ripk3*^*-/-*^ mice also exhibited a 47% decrease in their mean peripheral eosinophil population at 9 months when compared to wild-type littermate controls **(Figure 3E)**.

*Mlkl*^*-/-*^ and *Ripk3*^*-/-*^ mice of both sexes displayed genotype-dependent differences in non-white blood cell hematological parameters. *Mlkl*^*-/-*^ male mice showed significant increases in peripheral red blood cells (RBC) at 6 months when compared to wild-type littermate controls. Specifically, a 13% increase in the average absolute number and a 12% increase in the average ratio of RBC to the total volume of blood (percentage hematocrit) were observed at 6 months in comparison to littermate controls **(Supplementary Figure 3A, B)**. Fourteen-month-old *Mlkl*^*-/-*^ male mice also showed a 7% increase in the average number of RBC when compared to wild-type littermate controls, however, this was not reflected in percentage hematocrit **(Supplementary Figure 3A, B)**. Both *Mlkl*^*-/-*^ and *Ripk3*^*-/-*^ mice displayed significant age-dependent variations in the hemoglobin potential of circulating RBC compared to wild-type littermate controls. At 9 months, *Ripk3*^*-/-*^ males had a 5% increase in the mean corpuscular hemoglobin (MCH) measurements, whilst at 14 months, females exhibited a 44% decrease in MCH measurements, both compared with age-appropriate wild-type littermate controls **(Supplementary Figure 3C)**. Contrastingly, *Mlkl*^*-/-*^ male and female mice had significant decreases in MCH measurements compared to wild-type counterparts, with the former by 9% at 12 months and the latter by 3% at 6 months **(Supplementary Figure 3C)**. Only *Mlkl*^*-/-*^ mice displayed significant age-dependent differences in the size of the RBC. Whilst *Mlkl*^*-/-*^ males showed a 2% increase in MCV at 3 months, 6- and 9-month-old *Mlkl*^*-/-*^ females showed a 2% and 4% decrease in MCV, all compared to age-matched wild-type littermate controls **(Supplementary Figure 3D)**.

Finally, both *Mlkl*^*-/-*^ and *Ripk3*^*-/-*^ mice displayed significant age-dependent differences in platelet parameters. By 12 months of age *Mlkl*^*-/-*^ females exhibited a 26% increase in the average number of peripheral platelets when compared to wild-type littermate controls **(Supplementary Figure 3E)**. When compared to wild-type littermate controls, 14-month-old *Mlkl*^*-/-*^ females had a 10% decrease in mean platelet volume (MPV) and a corresponding 7% increase in mean platelet component (MPC) **(Supplementary Figure 3F, G)**. At 12 and 14 months of age, *Mlkl*^*-/-*^ male mice had a 16% increase and 16% decrease respectively in the MPV compared to wild-type littermate controls **(Supplementary Figure 3F)**. Consistent with these MPV findings, 12- and 14-month-old male *Mlkl*^*-/-*^ mice also exhibited an 11% decrease and a 16% increase in MPC compared to wild-type littermate controls **(Supplementary Figure 3G)**. The observed variation in peripheral lymphocyte counts and other haematological parameters appear to be tolerated at steady state. However, whether these differences may be disease-causing during challenges that reflect an everyday human scenario, such as surgical blood loss, pregnancy, or chemotherapy ablation, remains to be investigated.

### Bone marrow and splenic lymphocyte populations in female *Mlkl*^*-/-*^ mice are comparable to wild-type littermate controls

There are many possible causes for the elevated number of circulating lymphocytes observed in *Mlkl*^*-/-*^ mice, including mobilisation from lymphoid organs, enhanced generation, enhanced lifespan, or reduced clearance. Splenic and bone marrow lymphocytes numbers, specifically – CD4^+^ T cells, CD8^+^ T cells and CD19^+^ B cells – in 12-month-old female *Mlkl*^*-/-*^ showed no significant differences from wild-type littermate controls **(Figure 4A, B)**. Myeloid cell numbers, as well as lymphocyte numbers in the spleen and bone marrow were also comparable across genotypes at 12 months **(Figure 4A, B)**. Quantification of lymphocyte populations in the bone marrow and spleen did not provide insights as to the cause of increased peripheral lymphocyte numbers in female *Mlkl*^*-/-*^ mice, therefore histological examination was performed for all lymphoid organs at an age before the onset of lymphocytosis. Both *Mlkl*^*-/-*^ and *Ripk3*^*-/-*^ mice displayed typical lymphoid organs that resembled their respective wild-type littermate controls **(Figure 4C, Supplementary File 1)**.

**Figure 4.**
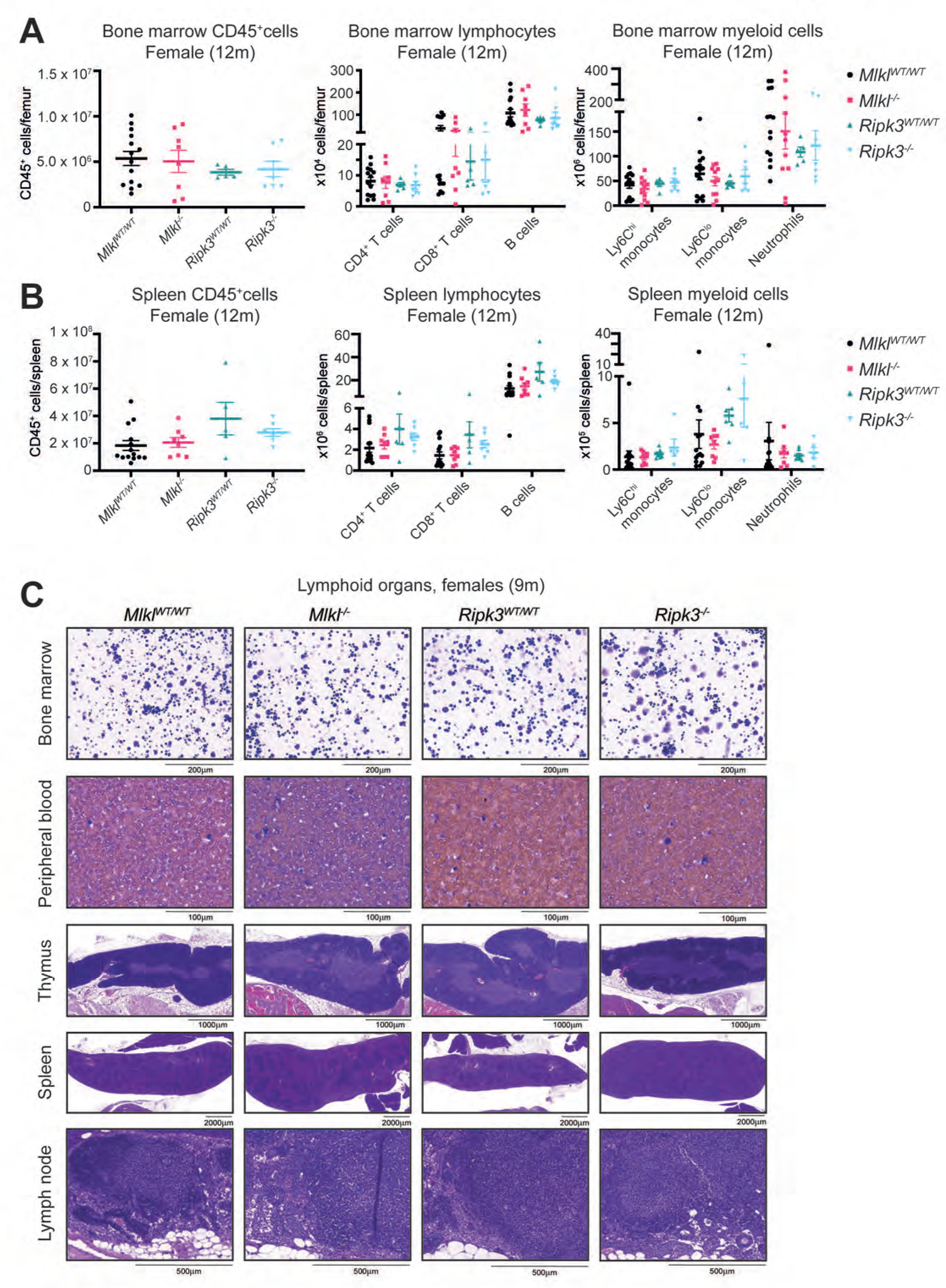
Female *Mlkl*^*-/-*^ mice have equivalent lymphocyte numbers in the spleen and bone marrow at 12 months of age. Quantification of CD45^+^ adaptive (CD4^+^ T cells, CD8^+^ T cells and B cells) and innate (Ly6C^hi^ monocytes, Ly6C^lo^ monocytes and neutrophils) immune cells in the bone marrow **(A)** and spleen **(B)** of 12-month-old female *Mlkl*^*-/-*^, *Ripk3*^*-/-*^ and wild-type littermate control mice. **(C)** H&E-stained sections of the lymphoid organs, bone marrow, peripheral blood, thymus, spleen, and lymph nodes, in female *Mlkl*^*-/-*^, *Ripk3*^*-/-*^ and wild-type littermate control mice at 9 months of age. Sections are representative of *n* = 2 – 6 mice per genotype. Each symbol represents a datum from one individual mouse and error bars represent mean ± SEM for *n* = 7 – 12. \**p* < 0.05 and \*\**p* < 0.01 were calculated using an unpaired, two-tailed Student’s t-test.

Male *Mlkl*^*-/-*^ mice also exhibited peripheral immune cell differences at 9 months of age, with a 62% reduction in the average circulating neutrophil number observed when compared to wild-type littermate controls **(Figure 3C)**. Primary and secondary lymphoid organs were assessed for neutrophil numbers at 6 months to understand the cause of this increase. Both male and female mice of all genotypes exhibited no significant differences in neutrophil or other lymphocyte and myeloid cell counts in the bone marrow, spleen, and inguinal lymph nodes **(Supplementary Figure 4A – F)**.

### Female *Mlkl*^*-/-*^ mice at 17 months of age exhibit a reduced number of inflammatory aggregates in connective tissue and muscle compared to wild-type littermate controls

*Mlkl*^*-/-*^ mice exhibit a delayed age-driven reduction in circulating lymphocytes and so we questioned whether the absence of MLKL protects against other phenotypic hallmarks of aging. Veterinary histopathologists, blind to genotype, examined the peripheral tissues of 9- and 17-month-old mice for evidence of inflammatory aggregates, a common finding in aging mice^52^ **(Figure 5A, Supplementary File 1 and 2)**. At 9 months of age, male and female *Mlkl*^*-/-*^ mice exhibited a similar degree of chronic sterile inflammation to wild-type littermate controls **(Figure 5B, Supplementary Figure 5A)**. It was noted, however, that 9-month-old female *Mlkl*^-/-^ mice exhibited an increasing trend in the total number of inflammatory foci in areas of connective tissue and muscle compared to wild-type littermate controls **(Figure 5C, Supplementary Figure 5B)**. These sites consist of the head (excluding glandular and nervous tissue), skin, hindleg, tail and sternum **(Supplementary Figure 5C – G)**. By 17 months, however, female *Mlkl*^-/-^ mice are protected against an age-dependent increase in sterile inflammation in connective tissue and muscle (**Figure 5A, C, Supplementary Figure 5C – G**). Compared to wild-type littermate controls, 17-month-old female *Mlkl*^-/-^ mice exhibit a 2.6-fold reduction in the number of inflammatory foci observed in the combined connective tissue and muscle (**Figure 5C**). This reduced number of inflammatory foci is not accompanied by any differences in the peripheral blood white blood cell populations, with 17-month-old *Mlkl*^-/-^ females and wild-type littermate controls exhibiting comparable numbers of peripheral lymphocytes, neutrophils, monocytes, and eosinophils **(Supplementary Figure 5H)**.

**Figure 5.**
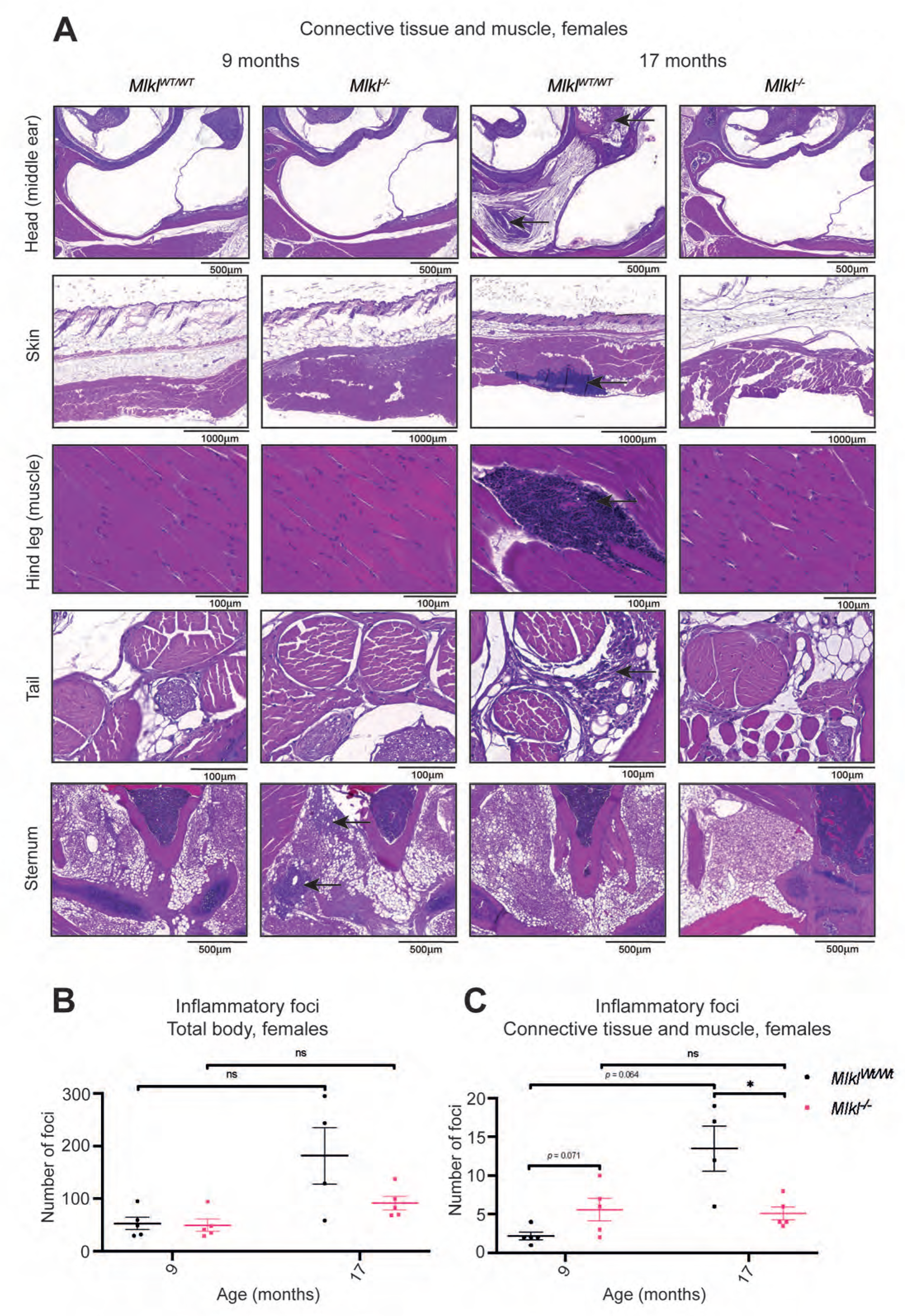
Female *Mlkl*^*-/-*^ mice are protected against chronic multifocal inflammation in connective and muscular tissue at 17 months of age. **(A)** Selected H&E sections of the head, skin, hind leg, tail, and sternum of 9- and 17-month-old female *Mlkl*^*WT/WT*^ and *Mlkl*^*-/-*^ mice. Inflammation is indicated by black arrows and H&E-stained sections are representative of *n* = 4 – 5 mice per genotype. **(B)** The total number of inflammatory foci identified in H&E-stained sections of all 44 sites across the body from female *Mlkl*^*WT/WT*^ and *Mlkl*^*-/-*^ mice. **(C)** The combined number of inflammatory foci identified in H&E-stained sections of the connective tissue and muscle that includes the head (excluding glandular and nervous tissue), hind leg, tail, skin, and sternum from female *Mlkl*^*WT/WT*^ and *Mlkl*^*-/-*^ mice. Each symbol represents a datum from one individual mouse and error bars display mean ± SEM for *n* = 4 – 5 mice. \**p* < 0.05 calculated using the non-parametric Mann-Whitney U test.

With exception of the connective tissue and muscle, blinded examination of inflammatory foci numbers across 44 sites revealed no significant differences between genotypes or sex at 9 and 17 months (**Supplementary File 1 and 2)**. Specifically, in tissues previously identified in MLKL-driven disease, such as the spinal cord, salivary glands, pancreas, liver, and kidney, inflammation was comparable irrespective of genotype and sex **(Supplementary Figure 5I – M, Supplementary File 1 and 2)**.

## DISCUSSION

Our extensive histopathological and immunophenotypic cohort analyses have identified several unique, sex-specific, differences between congenic C57BL/6J *Mlkl*^*-/-*^ and *Ripk3*^-/-^ mice and their wild-type littermates that emerge with age **(Figure 6)**. When measured as a percentage of body weight, male *Ripk3*^*-/-*^ mice had on average smaller, but histopathologically indistinguishable, spleens relative to wild-type littermate controls at 12 months. Several statistically significant findings were observed in hematological parameters across age in both *Mlkl*^*-/-*^ and *Ripk3*^-/-^ mice compared to wild-type littermate controls. Many of these parameters remained within normal range despite statistical significance, suggesting they are unlikely to assert biologically critical roles^53, 54, 55^. Of note, *Mlkl*^*-/-*^ mice exhibit increased circulating lymphocyte numbers relative to wild-type littermates at 12-14 months of age. A comprehensive and unbiased blind scoring of inflammatory foci in more than 44 different sites revealed a 2.6-fold decrease in background, sterile inflammation of the connective and muscular tissue in 17-month-old female *Mlkl*^*-/-*^ relative to age-matched wild-type littermates. It is important to consider, however, that these differences in age-related circulating lymphocyte numbers and tissue inflammation did not manifest in any overt differences in the general condition, mobility, or mortality of these mice up to 17 months of age.

**Figure 6.**
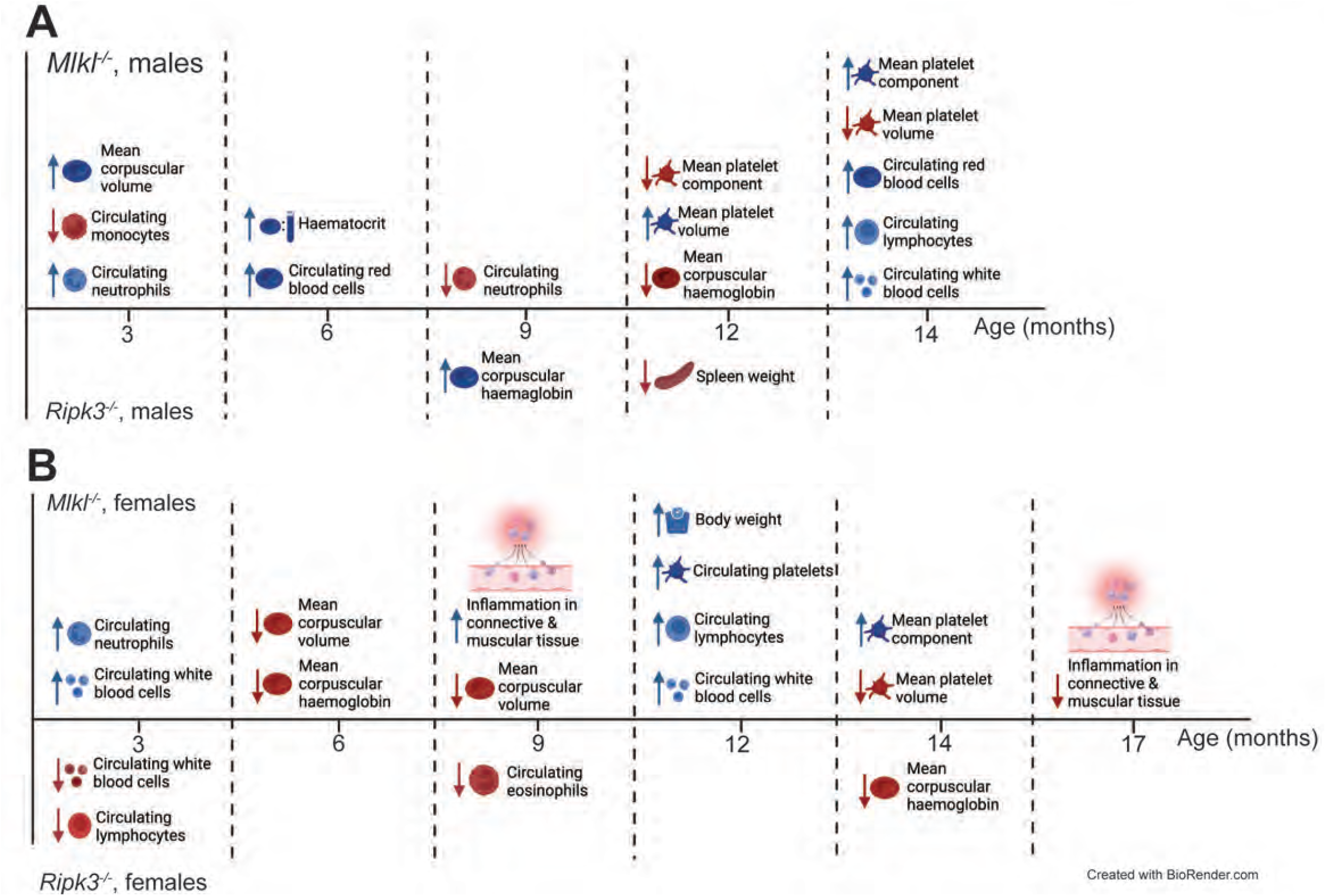
Summary of the identified phenotypic differences in *Mlkl*^*-/-*^ and *Ripk3*^*-/-*^ mice compared with age and sex-matched wild-type littermate controls. **(A)** Male *Mlkl*^*-/-*^ mice exhibit differences in circulating blood cell parameters at 3, 6, 9, 12 and 14 months of age. *Ripk3*^*-/-*^ male mice exhibit differences in circulating blood cell numbers at 9 and 12 months and a decrease in spleen weight at the latter age. **(B)** Both *Mlkl*^*-/-*^ and *Ripk3*^*-/-*^ females displayed phenotypes in the circulating blood at 3, 9 and 12 months, with the former also exhibiting significant differences at 6 months and the latter at 14 months. *Mlkl*^*-/-*^ females exhibit a significant decrease in the number of inflammatory foci identified in connective and muscular tissue at 17 months of age.

‘Inflammaging’ is the chronic increase in basal sterile inflammation that is known to underpin a large range of age-related disease^56, 57^. Whilst the processes of inflammaging are not completely understood, mouse models that undergo accelerated aging, such as NF-κB genetic knockouts, have provided clues to potential modulators and pathways^58, 59^. Furthermore, interventions in mice and humans intimately link anti-inflammatory effects and improved life span^60, 61, 62^. The significant reduction in the number of low-level inflammatory foci in the connective muscle and tissue of 17-month-old *Mlkl*^*-/-*^ females suggests that removal of necroptosis, or another non-necroptotic role of MLKL, limits age-dependent sterile inflammation in C57BL/6J mice. Inflammaging is known to be closely linked with immunosenescence, the age-dependent decline in immune function^57, 63^. The observed increase in the numbers of circulating lymphocytes at 12 and 14 months in female and male *Mlkl*^*-/-*^ mice, respectively, could represent a delay in the normal, age-dependent reduction in circulating lymphocytes in C57BL/6J mice^55^. Our observations of delayed immunosenescence and protection against inflammaging are suggestive that MLKL absence may contribute to widespread resistance to aging.

In the past four years, three previous reports have implicated human *MLKL* loss-of-function mutations in neurodegenerative or diabetic disease^46, 47, 48^. We did not detect evidence of either condition in appropriately aged *Mlkl*^*-/-*^ and *Ripk3*^*-/-*^ mice in this study. One such reason for this could be the approaches used for screening of disease pathologies, given there are more specific and accurate tests that could have been employed to identify nuanced phenotypes^50, 64, 65^. Nevertheless, we note that syndromes akin to the reports of severe human disease in patients with *MLKL* substitutions would have been detected in the current investigation. More likely, these human reports capture *MLKL* deficiency ‘in the real world’, where mutations occur alongside other genetic and/or environmental insults that are not consistent between individuals in the way that they are in laboratory mice. For example, in the two patients suffering from neurodegeneration, the *MLKL* loss-of-function mutation (*p*.*Asp369GlufsTer22*) also segregated with homozygous mutations in the *FA2H* and *ASP1S2* genes, which have each been linked to severe neurological syndromes^66, 67, 68^. Similarly, in the case of MODY diabetes, the heterozygous *MLKL* loss-of-function mutation also segregated with heterozygous deleterious mutations in the known MODY gene *PDX1*, as well as *ERN2, NIPAL4* and *SPTBN4* in affected individuals^48^. Whilst our investigations of *Mlkl*^*-/-*^ mice did not unveil any neurodegenerative or diabetic disease, it would be of interest to identify whether *MLKL* loss-of-function carriers presented with the phenotypes identified in our mice. Specifically, none of the three clinical reports provided information on the peripheral blood parameters of patients and it would be informative to know if, consistent with *Mlkl* knockout mice, they exhibit increased peripheral lymphocyte numbers. While it is likely that genetic context is critical to reveal patient phenotypes, whether our observed age-related changes in metabolic, hematological, and inflammatory parameters may act as modifiers of disease in the presence of physiological challenge is yet to be established.

Therapeutic targeting of the necroptotic signaling pathway is of immense clinical interest, exemplified by the active clinical trials of RIPK1-targeted small molecule compounds^43, 44, 45^. Despite no current first-in-human trials for RIPK3 or MLKL targeted drugs, their integral role in necroptosis makes them highly desirable potential targets (clinicaltrials.gov, accessed on 13 Sep 2022). The age-dependent phenotypes that we observed in *Mlkl*^*-/-*^ mice highlight that long-term inhibition may have several important age-related health benefits. However, differing age-related phenotypes in *Ripk3*^*-/-*^ and *Mlkl*^*-/-*^ mice suggest there are advantages to therapeutically targeting RIPK3 or MLKL depending on the disease context. Our study provides provocative evidence of age-related MLKL and RIPK3 roles that will be of immense interest as anti-necroptotic inhibitors advance toward the clinic.

## METHODS

### Animal ethics

All mice used in this study were housed at a temperature and humidity-controlled facility at the Walter and Eliza Hall Institute of Medical Research. Operating on a 12h:12h day-night cycle, this is a specific pathogen-free facility. All experiments were approved by the WEHI Animal Ethics Committee following the Prevention of Cruelty to Animals Act (1996) and the Australian National Health and Medical Research Council Code of Practice for the Care and Use of Animals for Scientific Purposes (1997).

### Mice

The generation of the *Mlkl*^*-/-*^ mice used in this study was reported previously^5^. The *Ripk3*^*-/-*^ mice were generated by the MAGEC laboratory (WEHI) on a C57BL/6J background. To generate the *Ripk3*^*-/-*^mice, 20 ng/μl of Cas9 mRNA, and 10 ng/μl of sgRNAs (AACTTGACAGAAGACATCGT and GATTCTCTGAAGTCTACTTG; targeting the 5’ UTR-Exon1 and Exon 10-3’ UTR junctions, respectively, for deletion of the intervening locus) were injected into the cytoplasm of fertilized one-cell stage embryos generated from wild-type C57BL/6J breeders. Twenty-four hours later, two-cell stage embryos were transferred into the uteri of pseudo-pregnant female mice. Viable offspring were genotyped by next-generation sequencing. Targeted animals were backcrossed twice to wild-type C57BL/6J to eliminate off-target mutations. Mice aged to 3 months were between 13 – 16 weeks, 6 months between 26 – 30 weeks, 9 months between 35 – 43 weeks, 12 months between 49 – 60 weeks except for ADVIA data where mice were aged between 49 – 58 weeks, 14 months between 60 – 64 weeks of age and 17 months between 73 – 75 weeks. Mice were checked twice weekly until 52 weeks old, after which they were checked three times weekly. All mice in this study were housed within the same facility, with genetic knockout mice housed in the same room as their wild-type littermate controls. *Mlkl*^*-/-*^ and *Ripk3*^*-/-*^ colonies were maintained in different rooms within the same facility.

### Hematological analysis

Cardiac or sub-mandibular blood from mice aged between 3 – 17 months was collected into EDTA-coated tubes. Blood was diluted between 2- and 10-fold in DPBS for automated blood cell quantification using an ADVIA 2120i hematological analyzer on the same day as harvest. Accu-Chek Performa Blood Glucose Meter measured blood glucose on undiluted blood within two hours of harvest. Any mice that had been observed fighting and exhibited overt skin wounds resulting from fighting were excluded from hematological analyses.

### Whole body histopathology, 9 and 17 months of age

A comprehensive histological examination of 9-month-old male and female *Mlkl*^*WT/WT*^, *Mlkl*^*-/-*^, *Ripk3*^*WT/WT*^, and *Ripk3*^*-/-*^ mice and 17-month-old female *Mlkl*^*WT/WT*^ and *Mlkl*^*–/–*^ mice was completed by histopathologists at Phenomics Australia, Melbourne. 9-month-old mice were dispatched to Phenomics Australia in – separate cohorts, whilst 17-month-old female mice were all dispatched in one singular cohort on the same day. Mice were euthanized by CO_2_ inhalation. All soft tissues, except for testes, epididymes, eyes and Harderian glands, were placed into 10% v/v neutral buffered formalin (NBF) for 24 hours before being transferred into 10% v/v NBF. Testes and epididymes were placed into Bouin’s fixative and the eyes and Harderian glands into Davidson’s fixative. Skeletal tissues were fixed for 48 hours in 10% v/v NBF, followed by 48 hours of 10% v/v formic acid for decalcification and then back into 10% v/v NBF for a further 24 hours. Following fixation, tissues were moved for automated processing to paraffin using the Sakura VIP6 auto processor. Paraffin blocks of embedded tissues were sectioned at 5 µm thickness, with one representative section taken through the entire tissue. Sections were stained with hematoxylin and eosin on the Leica Autostainer XL/Leica CV5030 coverslipper. X-rays were obtained using the portable X-ray system iRayD3 (DX-3000L).

Trained histopathologists examined all tissues from mice at 9 or 17 months of age for inflammation, quantifying the number of inflammatory foci in each section, blinded to genotype. The size of the brain was quantified by measuring the length, width, and height. Senior Veterinary Pathologist, Prof. John W. Finnie from the School of Medicine, University of Adelaide performed a detailed neuropathological examination on all mice and confirmed the full body findings for all 9-month-old mice. Senior Veterinary Pathologist, Dr Lorna Rasmussen confirmed the full body findings for all 17-month-old mice.

### Organ tissue harvest and processing, 12 months of age

Twelve-month-old mice were euthanized by CO_2_ inhalation immediately before organ dissection. Each organ was weighed before fixation in 10% v/v neutral buffered formalin and long-term storage in 80% v/v ethanol. Mice that had been observed fighting and exhibited overt skin wounds resulting from fighting were plotted as hollowed-out symbols. H&E staining of harvested spleen was completed by the WEHI Histology Facility. Heart, colon, ileum, spleen, testes, and liver organ samples taken from 12-month-old mice were lysed in ice-cold RIPA buffer supplemented with Protease & Phosphatase Inhibitor Cocktail (Cell Signaling Technology 5872 S, 50 mg/ml) and Benzonase (Sigma-Aldrich E1014, 100 U/mL). Tissue lysates were homogenized using the Qiagen TissueLyser II (30 Hz, 1 min). Proteins were harvested in 2 x SDS-loading buffer and visualised as per the western blot protocol below.

### Flow cytometry

To analyze immune cells (innate and adaptive) in the spleen, bone marrow or inguinal lymph nodes of 6- and 12-month-old *Mlkl*^*WT/WT*^, *Mlkl*^*-/-*^, *Ripk3*^*WT/WT*^, and *Ripk3*^*-/-*^ mice, single-cell suspensions were subjected to osmotic red blood cell lysis and incubated with a combination of antibodies: CD4-BV421 (BD Biosciences #562891), CD8-PeCy7 (BD Biosciences #561097), CD19-PerCPCy5.5 (BD Biosciences #551001), CD11b-BV510 (BD Biosciences #562950) or CD11b-BV786 (BD Biosciences #740861), CD45-Alexa700 (BD Biosciences #560566), Ly6G-V450 (BD Biosciences #560603) and Ly6C-APCCy7 (BD Biosciences #560596). Samples were analyzed on an Aurora Cytex flow cytometer, with the automated volume calculator used to quantify absolute cell numbers. FlowJo v10.8 was used for all analyses.

### Cell line generation and maintenance

Primary mouse dermal fibroblasts (MDFs) were prepared from the skin taken from the tails of 6-week-old *Ripk3*^*WT/WT*^ or *Ripk3*^*–/–*^ mice. Primary MDFs were immortalized by stable lentiviral transduction with a DNA construct encoding the SV40 large T antigen. MDFs were cultured in Dulbecco’s Modified Eagle Medium (DMEM) + 8% (v/v) fetal calf serum (FCS) at 37 °C with 10% CO_2_. Cell lines were routinely monitored for mycoplasma.

### IncuCyte cell death assays

MDFs were seeded into 96-well plates at 8 × 10^3^ cells/well and left to settle for 6 hours. The next day MDFs were stimulated in Phenol Red-free DMEM (ThermoFisher Scientific) supplemented with 2% FCS, 1mM L-GlutMAX, G/P/S, 1mM Na pyruvate, SYTOX Green (5µM; Invitrogen S7020) and DRAQ5 (10µM; Thermofisher #62251). MDFs were stimulated with combination treatments of TNF (100 ng/mL), Smac-mimetic compound A (500 nM; TS) or TS plus pan-caspase inhibitor IDN-6556 (5 µM; TSI), in the absence or presence of RIPK3 inhibitor GSK’872 (1 µM). Cells were moved into a SX5 system (37 °C, 10% CO_2_; Essen Bioscience) and imaged with a 10x objective every hour. Percentage cell death values were quantified by the number of dead cells (SYTOX Green-positive) out of the total live cell number (DRAQ5 positive) using the IncuCyte v.2022A software.

### Western blot

Immortalized MDFs were plated at 3 × 10^4^ cells/well and left to adhere overnight. The next day, cells were stimulated as indicated with a combination of treatments TNF (100 ng/mL), Smac-mimetic Compound A (500 nM), pan-caspase inhibitor IDN-6556 (5 µM), and GSK-872 (1 µM) for 4 hours. All cells were harvested in 2 x SDS Laemmli lysis buffer and boiled at 100 °C for 10-15 minutes. Proteins were resolved by 4-15% Tris-Glycine gel (Bio-Rad) and transferred to nitrocellulose membrane, before probing with indicated antibodies. Uncropped blots are shown as supplemental data.

### Reagents

Antibodies: Rat anti-mMLKL 5A6 and rat anti-mRIPK3 1H12 were produced in-house^69^ (5A6 available from Merck as MABC1634). Mouse anti-actin (A-1978) was purchased from Sigma-Aldrich, rabbit anti-mouse pMLKL (D6E3G) was purchased from CST and rabbit anti-mouse pRIPK3 (GEN135-35-9) was kindly provided by Genentech^70^. Recombinant hTNF-Fc (produced in-house) and the Smac mimetic, Compound A, have been previously described^71, 72^. Pan-caspase inhibitor IDN-6556 was provided by Tetralogic Pharmaceuticals and GSK’872 was sourced from SynKinase.

### Statistical analysis

All data points signify independent experimental repeats, and/or biologically independent repeats. All *p* values were calculated in Prism using an unpaired, two-tailed Student’s t-test or non-parametric Mann-Whitney U test as indicated. Asterisks signify that *p* ≤ 0.05 (*), *p* ≤ 0.01 (**), *p* ≤ 0.001 (***).

## ACKNOWLEDGEMENTS

We thank the following people for their technical assistance with the experiments completed in this manuscript; Aira Nuguid and Tina Cardamone (Phenomics Australia Histopathology and Slide Scanning Service-The University of Melbourne), WEHI Bioservices, WEHI Histology Centre and WEHI Cytometry Facility. The generation of the *Ripk3*^*-/-*^ mice used in this study was supported by Phenomics Australia and the Australian Government through the National Collaborative Research Infrastructure Strategy (NCRIS) program. We thank Katherine Martin for the provision of important resources and expertise. We are grateful to the National Health and Medical Research Council for fellowship (J.M.H., 1142669; A.L.S., 2002965; J.M.M., 1172929), grant (J.M.H., 2011584) and infrastructure (IRIISS 9000719); and the Victorian State Government Operational Infrastructure Support scheme. We also acknowledge scholarship support for S.E.G (Australian Government Research Training Program Stipend Scholarship), S.E.G. (Wendy Dowsett Scholarship), S.C. (Walter and Eliza Hall Handman PhD Scholarship) and E.C.C.T. (Attracting and Retaining Clinician-Scientists’ Scholarship).

## CONTRIBUTIONS

Conceptualization: E.C.T, S.E.G., J.M.M., J.M.H. Methodology: E.C.T., S.E.G., J.D., H.A., S.C., C.H., K.M.P., P.G., A.L.S. Resources: A.H., A.J.K., U.N., J.M.M., J.M.H. Statistical oversight: C.S.L., A.L.G. Supervision: K.E.L., A.L.S., J.M.M., J.M.H. Funding acquisition: J.M.M., J.M.H. Writing: E.C.T., S.E.G., J.M.M. and J.M.H. co-wrote the paper with input from authors.

