## Supplementary Figures for "MLKL deficiency protects against low-grade, sterile inflammation in aged mice"

**Supplementary Figure 1. *Ripk3*$^{-/-}$ mice on a C57BL/6J background have no detectable RIPK3 expression and do not execute necroptotic cell death. (A)** sgRNA targeting strategy for the generation of RIPK3 deficient mice. **(B)** *Ripk3*$^{-/-}$ mice are born according to Hardy-Weinberg equilibrium as observed in the distribution of genotypes from *Ripk3*$^{+/-}$ heterozygous intercrosses. **(C)** Immunoblot for RIPK3 protein on extracts of tissues from three *Ripk3*$^{-/-}$ mice and one wild-type mouse. **(D)** Immortalised mouse dermal fibroblasts (MDFs) were isolated from wild-type and *Ripk3*$^{-/-}$ mice and stimulated for 18 hours for quantification of the percentage of SYTOX-Green positive cells using IncuCyte SX5 live cell imaging. One independent wild-type MDF cell line and three independent *Ripk3*$^{-/-}$ MDF cell lines were assayed in $n = 2$ experiments, with the mean depicted in solid line and each replicate displayed as dots. **(E)** Three *Ripk3*$^{-/-}$ independent MDF cell lines and one wild-type MDF cell line were stimulated as indicated for 4 hours for western blot analysis. **(F-H)** Absolute and relative weight of pancreas **(F)**, liver **(G)** and kidney **(H)** from 12-month-old *Mlkl*$^{-/-}$, *Ripk3*$^{-/-}$ and wild-type littermate control mice. Each symbol represents a datum from one individual mouse and error bars represent mean ± SEM for $n = 7 – 11$. Hollowed-out symbols represent data from fighting male mice and $*p < 0.05$ was calculated using an unpaired, two-tailed Student's t-test.

### Supplementary Figure 1

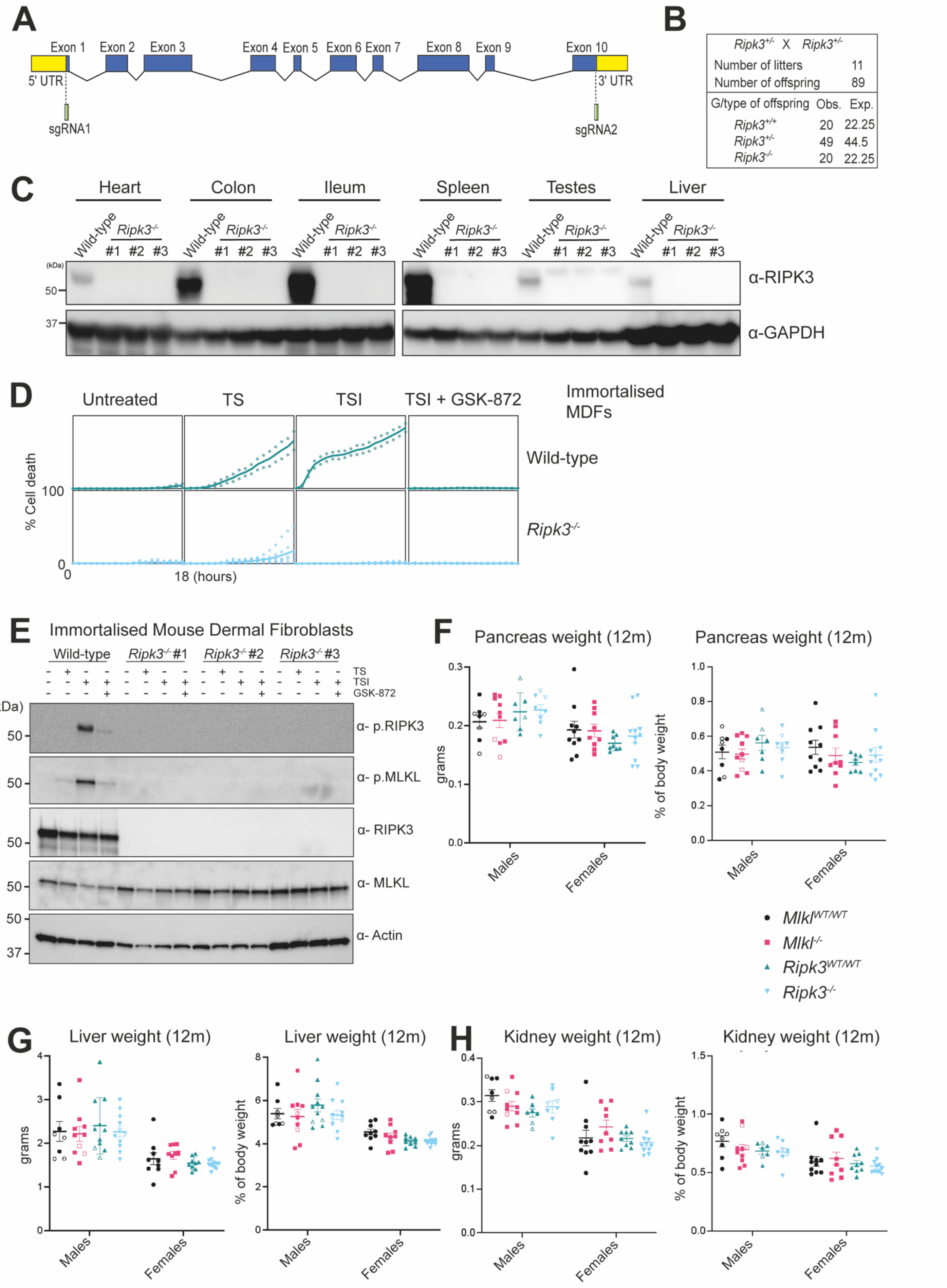

**Supplementary Figure 2. Nine-month-old *Mlkl*$^{-/-}$, *Ripk3*$^{-/-}$ and wild-type littermate control mice are ostensibly normal and neurologically intact. (A)** Macroscopic appearance of *Mlkl*$^{-/-}$, *Ripk3*$^{-/-}$ and littermate control male mice at 9 months of age. **(B)** H&E-stained sections of the levels I, II and III of the brain and **(C)** X-ray images of the head, torso, pelvis, and hind leg from 9-month-old female *Mlkl*$^{-/-}$, *Ripk3*$^{-/-}$ and wild-type littermate control mice. Level I includes the cortex, corpus callosum, caudate putamen and lateral ventricles; level II includes the hippocampus, thalamus, hypothalamus, and lateral and third ventricles, and level III, the cerebellum, pons and fourth ventricle. **(D)** Histopathologist quantification of brain size by length, width, and height at 9 months old in male and female *Mlkl*$^{-/-}$, *Ripk3*$^{-/-}$ and littermate control mice. Whole body images **(A),** H&E-stained sections **(B)** and X-ray images **(C)** are representative of $n = 2 – 5$ mice per genotype. Each symbol represents a datum from one individual mouse and error bars represent mean ± SEM for $n = 2 – 5$. Hollowed-out symbols represent data from fighting male mice.

### Supplementary Figure 2

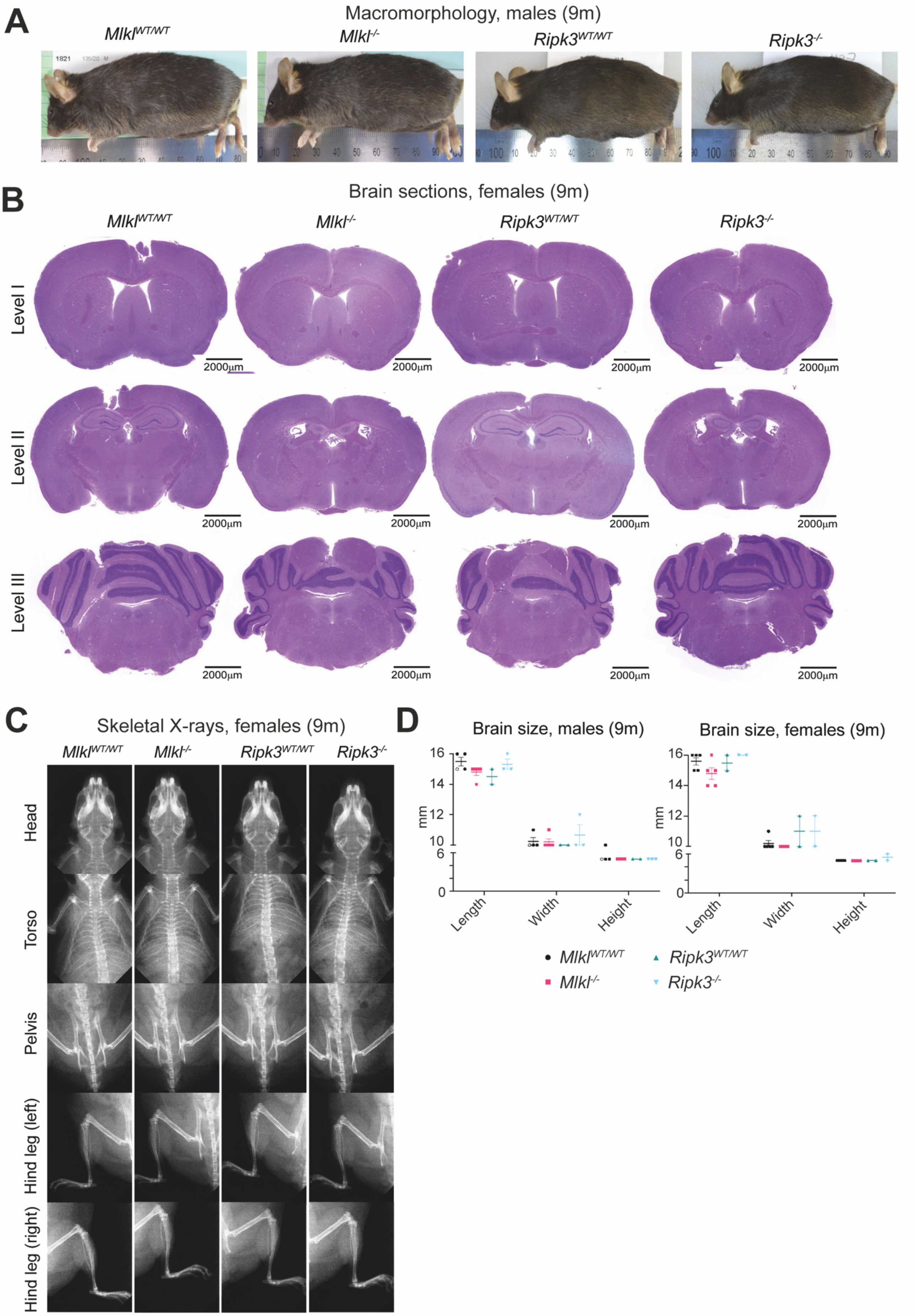

**Supplementary Figure 3. Female *Mlkl*$^{-/-}$ mice exhibit increased circulating platelet numbers at 12 months of age.** ADVIA hematology quantification of circulating red blood cells **(A)**, percentage hematocrit **(B)**, mean corpuscular hemoglobin **(C)**, mean corpuscular volume **(D)**, platelets **(E),** mean platelet volume **(F)** and mean platelet component **(G)** in sex separated *Mlkl*$^{-/-}$, *Ripk3*$^{-/-}$ and wild-type littermate controls at 3 – 14 months of age. Each symbol represents a datum from one individual mouse and error bars represent mean ± SEM for $n$= 2 – 22. *$p < 0.05$, **$p < 0.01$, ***$p < 0.001$ calculated using an unpaired, two-tailed Student's t-test.

### Supplementary Figure 3

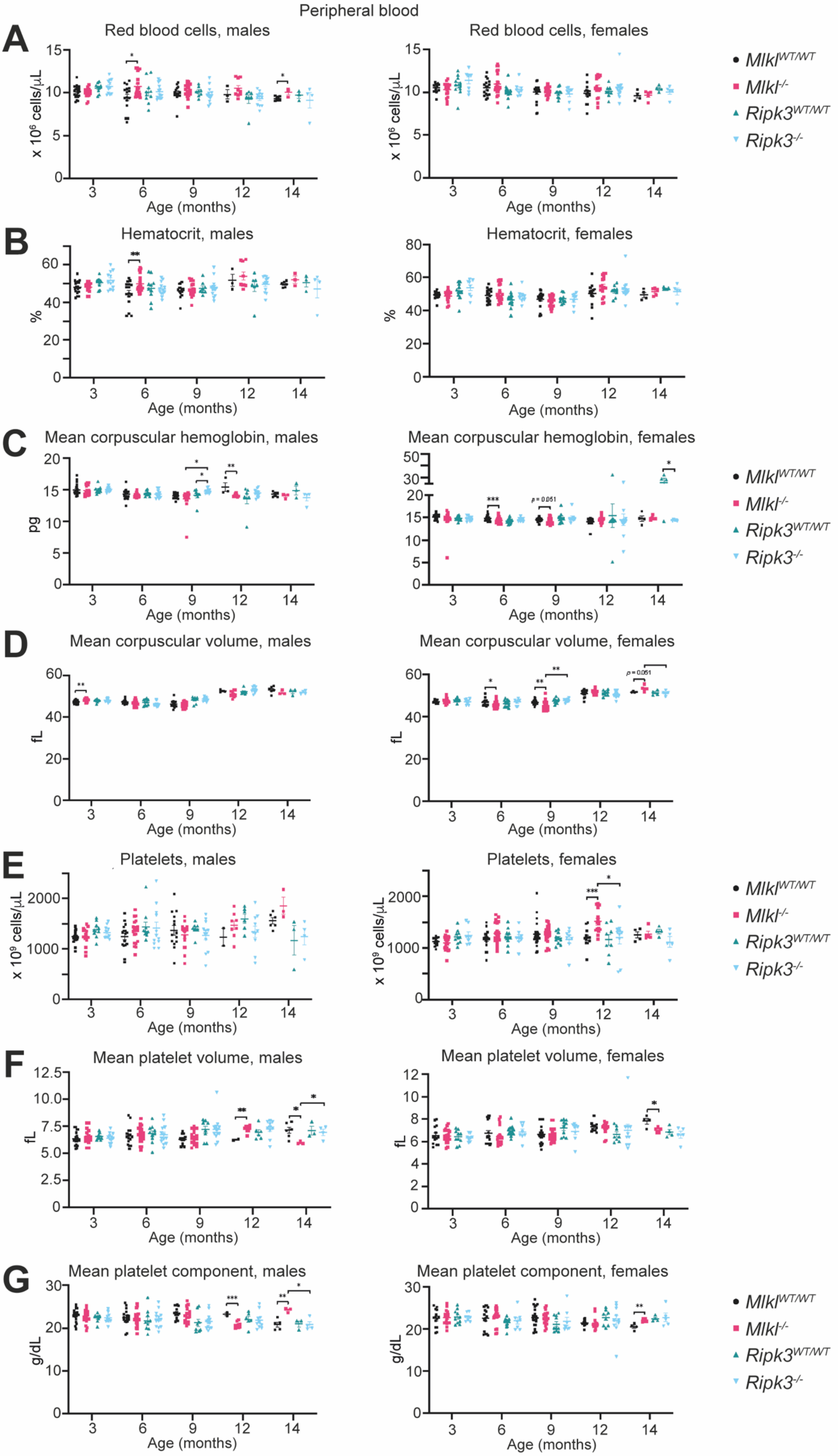

**Supplementary Figure 4. Six-month-old *Mlkl*$^{-/-}$ mice show comparable immune cell numbers in lymphoid organs compared to wild-type littermate controls. (A – F)** Quantification of adaptive (CD4$^{+}$ T cells, CD8$^{+}$ T cells and B cells) and innate (Ly6C$^{hi}$ monocytes, Ly6C$^{lo}$ monocytes and neutrophils) immune cells in the bone marrow, spleen, and inguinal lymph nodes of 6-month-old *Mlkl*$^{-/-}$ male and female mice compared to wild-type littermate controls. Each symbol represents a datum from one individual mouse and error bars represent mean ± SEM for $n$ = 8 – 11. Hollowed-out symbols represent data from fighting male mice.

### Supplementary Figure 4

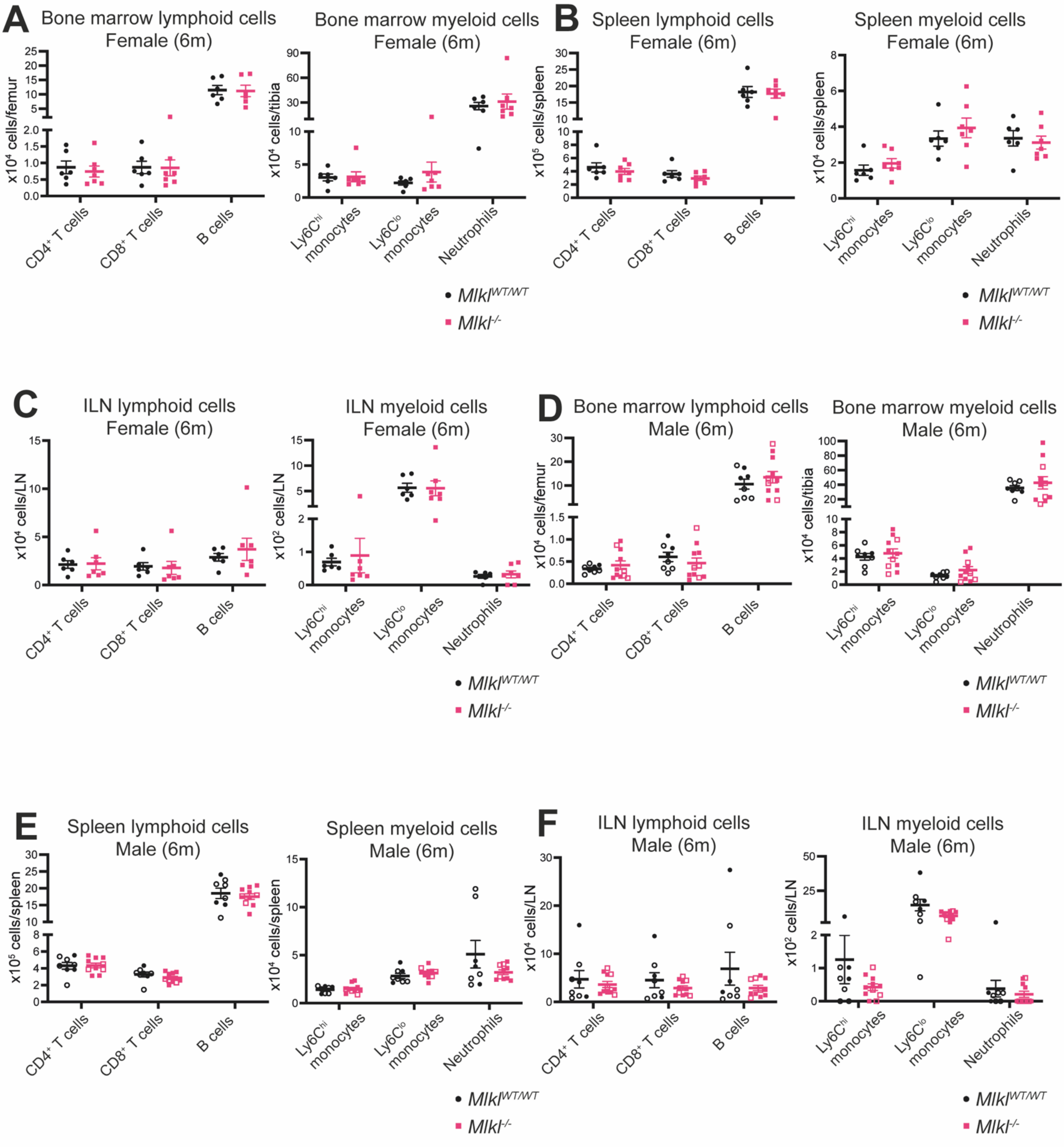

**Supplementary Figure 5. Female *Mlkl*$^{-/-}$ mice do not exhibit site-specific differences in inflammation compared to littermate controls at 9 months.** The total number of inflammatory foci identified in H&E-stained sections of all 44 sites across the body **(A)** or combined connective tissue and muscle **(B)** in 9-month-old male *Mlkl*$^{WT/WT}$ and *Mlkl*$^{-/-}$ mice. The number of inflammatory foci identified in sections of the head **(C)**, skin **(D)**, hind leg **(E)**, tail **(F)**, sternum **(G)**, spinal cord **(I),** salivary glands **(J)**, pancreas **(K)**, liver **(L)** and kidneys **(M)** from female *Mlkl*$^{WT/WT}$ and *Mlkl*$^{-/-}$ mice at 9 and 17 months. **(H)** ADVIA hematology quantification of peripheral white blood cell numbers in female *Mlkl*$^{WT/WT}$ and *Mlkl*$^{-/-}$ mice at 17 months old. Each symbol represents one individual mouse sampled and error bars represent mean ± SEM for $n = 3 – 5$. Hollowed-out symbols represent data from fighting male mice. *$p<0.05$ calculated using the non-parametric Mann-Whitney U test.

### Supplementary Figure 5

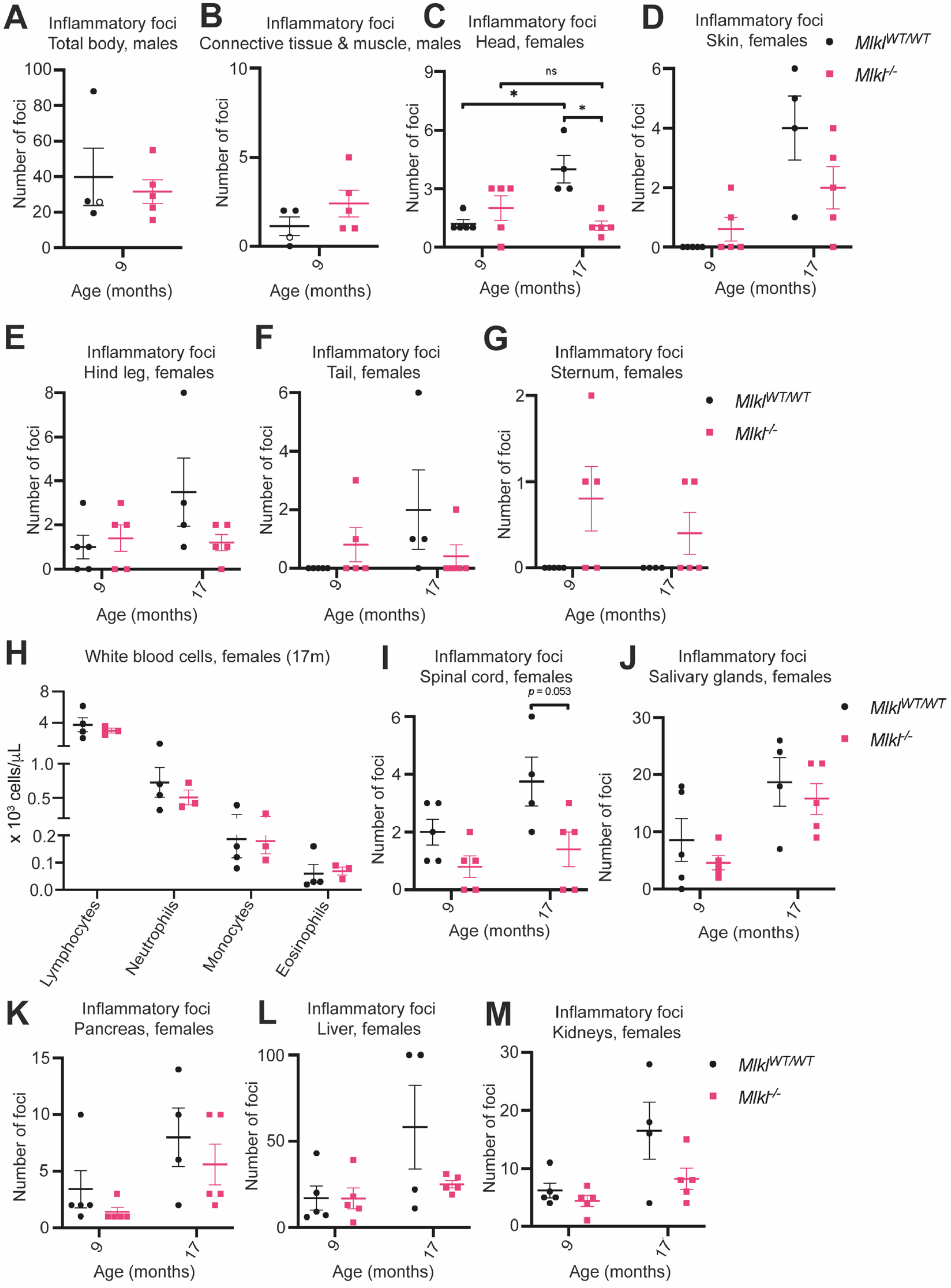

**Supplementary File 1. Macro- and microscopic observations of 9- and 17-month-old *Mlkl*$^{-/-}$ mice compared to wild-type littermate controls.** Macroscopic observations include gross neurological tests, body condition scoring, anthropometric measurements, and organ size. Microscopic observations include a detailed description of H&E-stained slides of 44 unique anatomical locations by professional histopathologists.

**Supplementary File 2. Blind semi-quantitative scoring of the number and size of inflammatory foci identified in 44 unique anatomical sites in 9- and 17-month-old *Mlkl*$^{-/-}$ mice compared to wild-type littermate controls.** The number of inflammatory foci is based on the average number of foci identified across all H&E-stained sections of the tissue/organ. A focus is defined as an aggregate of leukocyte cells > 10 in number. The size of inflammatory foci is based on the largest focus identified across all sections of the tissue/organ where 1 = < 50 cells per focus, 2 = > 50 cells per focus and 3 = bridging of foci. This analysis was blinded and performed by professional histopathologists.
